# Intronic enhancer region governs transcript-specific BDNF expression in neurons

**DOI:** 10.1101/2020.11.25.398156

**Authors:** Jürgen Tuvikene, Eli-Eelika Esvald, Annika Rähni, Kaie Uustalu, Anna Zhuravskaya, Annela Avarlaid, Eugene V. Makeyev, Tõnis Timmusk

## Abstract

Brain-derived neurotrophic factor (BDNF) controls the survival, growth, and function of neurons both during the development and in the adult nervous system. BDNF gene is transcribed from several distinct promoters generating transcripts with alternative 5’ exons. BDNF transcripts initiated at the first cluster of exons have been associated with the regulation of body weight and various aspects of social behavior, but the mechanisms driving the expression of these transcripts have remained poorly understood. Here, we identify an evolutionarily conserved intronic enhancer region inside the BDNF gene that regulates both basal and stimulus-dependent expression of the BDNF transcripts starting from the first cluster of 5’ exons in neurons. We further uncover a functional E-box element in the enhancer region, linking the expression of BDNF and various pro-neural basic helix-loop-helix transcription factors. Collectively, our results shed new light on the cell type- and stimulus-specific regulation of the important neurotrophic factor BDNF.

## INTRODUCTION

Brain-derived neurotrophic factor (BDNF) is a secreted protein of the neurotrophin family (Park and Poo, 2013). During the development, BDNF promotes the survival of various sensory neuron populations (Ernfors et al., 1994; Jones et al., 1994). In the adult organism, BDNF is also required for the proper maturation of synaptic connections and regulation of synaptic plasticity (Korte et al., 1995; Park and Poo, 2013). Defects in BDNF expression and signaling have been implicated in various neuropsychiatric and neurodegenerative diseases, including major depression, schizophrenia, Alzheimer’s disease and Huntington’s disease (Autry and Monteggia, 2012; Burbach et al., 2004; Jiang and Salton, 2013; Murray et al., 1994; Ray et al., 2014; Wong et al., 2010; Zuccato et al., 2001; Zuccato and Cattaneo, 2009).

Murine BDNF gene contains eight independently regulated non-coding 5’ exons (exons I-VIII) followed by a single protein-coding 3’ exon (exon IX). Splicing of one of the alternative exons I-VIII with the constitutive exon IX gives rise to different BDNF transcripts (Aid et al., 2007). Additionally, transcription can start from an intronic position upstream of the coding exon producing an unspliced 5’ extended variant of the coding exon (exon IXa-containing transcript) (Aid et al., 2007). The usage of multiple promoters enables complex cell-type and stimulus-specific BDNF expression (reviewed in West et al., 2014). For instance, BDNF exon I, II, and III-containing transcripts show mainly nervous system-specific expression patterns, whereas BDNF exon IV and VI-containing transcripts are expressed in both neural and non-neural tissues (Aid et al., 2007; Timmusk et al., 1993). Similar expression patterns for different BDNF transcripts are also observed in human (Pruunsild et al., 2007). Notably, different BDNF transcripts have distinct contribution to various aspects of neural circuit functions and behavior (Hallock et al., 2019; Hill et al., 2016; Maynard et al., 2016a, 2018a; McAllan et al., 2018; Sakata et al., 2009).

In addition to proximal promoter regions, the complex regulation of gene expression is often controlled by distal regulatory elements called enhancers (reviewed in Buecker and Wysocka, 2012). Enhancers are usually active in a tissue- and cell type-specific manner (reviewed in Heinz et al., 2015; Wu et al., 2014), and can be located inside or outside, upstream or downstream of the target gene, within another gene or even on a different chromosome (Banerji et al., 1981; Lettice et al., 2003; reviewed in Ong and Corces, 2011). Many enhancers are activated only after specific stimuli which cause enrichment of active enhancer-associated histone modifications and increased chromatin accessibility (Su et al., 2017). Genome-wide analysis has proposed approximately 12000 neuronal activity-regulated enhancers in cortical neurons (Kim et al., 2010). Importantly, dysregulation of enhancers or mutations in proteins that participate in the formation of enhancer-promoter complexes are associated with a variety of disorders, including neurodegenerative diseases (reviewed in Carullo and Day, 2019).

Previous studies from our laboratory have suggested that the expression of the BDNF gene is also regulated via distal regulatory regions. Notably, the induction of BDNF mRNA after BDNF-TrkB signaling in neurons seems to depend on unknown distal regulatory regions (Esvald et al., 2020). Furthermore, dopamine-induced expression of BDNF in astrocytes is controlled by an unknown regulatory region within the BDNF gene locus (Koppel et al., 2018). Here, we identify a novel enhancer region in the BDNF gene located downstream of the BDNF exon III and show that this regulatory element selectively activates basal and stimulus-dependent expression of the exon I, II and III-containing BDNF transcripts in neurons.

## RESULTS

### 1. BDNF +3 kb region shows enhancer-associated characteristics in mouse and human brain tissue

To uncover novel enhancer regions regulating BDNF expression in the central nervous system, we started with bioinformatic analysis of the enhancer-associated characteristics. Active enhancers are characterized by nucleosome-free DNA that is accessible to transcription factors and other DNA binding proteins. Chromatin at active enhancer regions typically has distinct histone modifications – H3K4me1, a hallmark of enhancer regions, and H3K27ac, usually associated with active regulatory regions. Active enhancers also bind RNA polymerase II and are bidirectionally transcribed from the regions marked by enhancer-associated histone modifications giving rise to non-coding enhancer RNAs (eRNAs) (Nord and West, 2020). Based on the mouse brain tissue ChIP-seq data from the ENCODE project and transcription start site (TSS) data from the FANTOM5 consortium, a region ~3 kb downstream of BDNF exon I TSS has prominent enhancer-associated features (Figure 1A). First, the +3 kb region is hypersensitive to DNaseI, indicative of an open chromatin structure. Second, ChIP-seq data shows that this region is enriched for H3K4me1, H3K4me3 and H3K27ac modifications. Third, the +3 kb region interacts with RNA polymerase II, with a strong evidence for bidirectional transcription according to the FANTOM5 CAGE database. Finally, the region is conserved between mammals, pointing at its possible functional importance.

**Figure 1.**
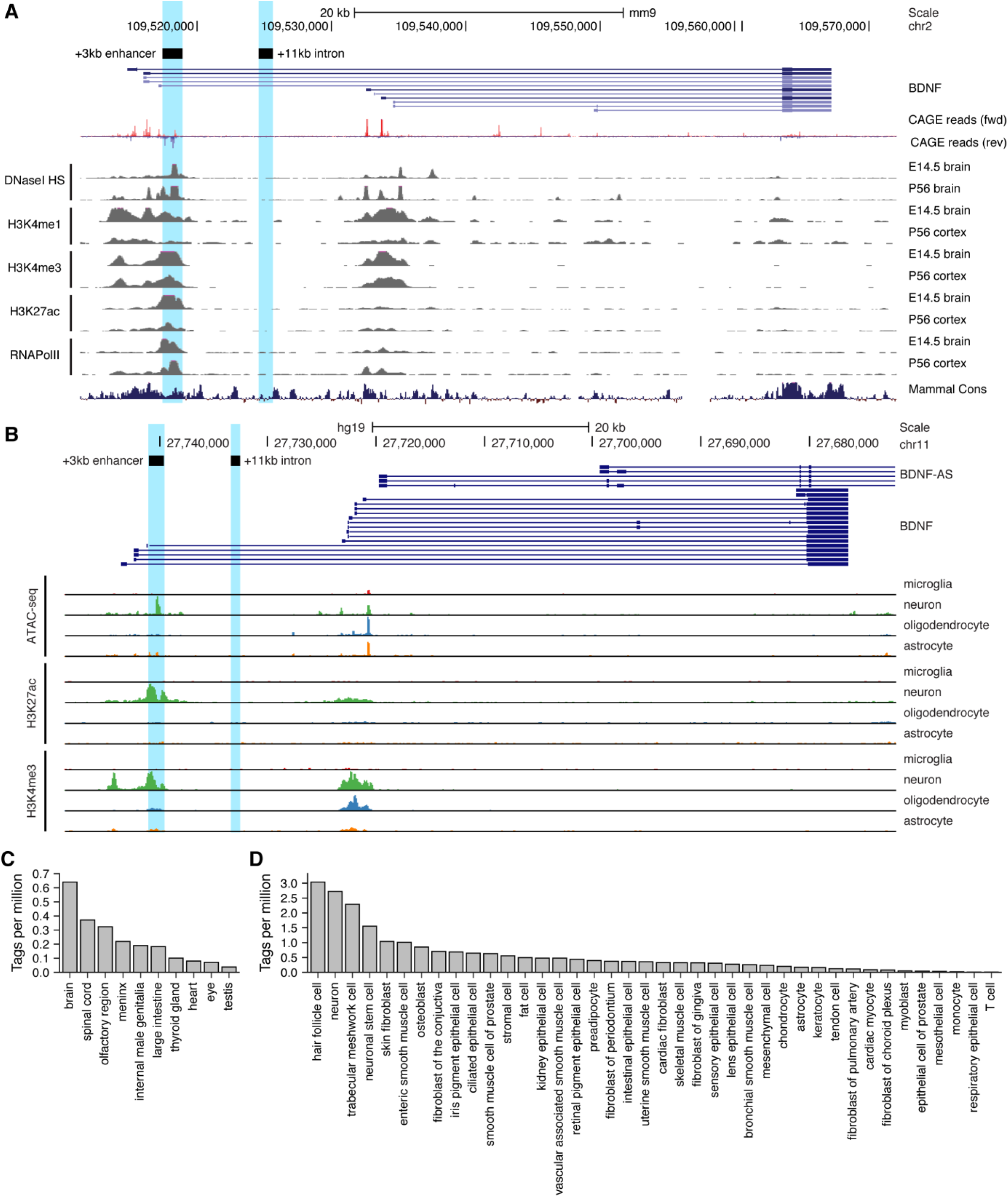
Region downstream of BDNF exon III shows enhancer-associated characteristics in mouse and human neural tissues. UCSC Genome browser was used to visualize (A) DNaseI hypersensitivity sites and ChIP-seq data from the ENCODE project in mouse brain tissue, CAGE data of transcription start sites from the FANTOM5 consortium (all tissues and cell types), and (B) open chromatin (ATAC-seq) and ChIP-seq in different human brain cell-types by (Nott et al., 2019). E indicates embryonic day, P postnatal day. Signal clipping outside the visualization range is indicated with purple color. The +3 kb region, a potential enhancer of the BDNF gene, and +11 kb intronic region, a negative control region used in the present study were converted from rat genome to mouse or human genome using UCSC Liftover tool and are shown as light blue. The names of the regions represent the distance of the respective region from rat BDNF exon I transcription start site. (C, D) +3 kb enhancer region (chr11:27693843-27694020, hg19 genome build) eRNA expression levels based on CAGE sequencing data from the FANTOM5 project obtained from the Slidebase tool (Ienasescu et al., 2016, http://slidebase.binf.ku.dk). eRNA expression levels were grouped by different tissue types (C) or different cell types (D). Only tissue and cell types with non-zero eRNA expression are shown.

We next used H3K27ac ChIP-seq data from Nord et al (Nord et al., 2013) to determine the activity of the potential enhancer region in different tissues throughout the mouse development. We found that the +3 kb region shows H3K27ac mark in the mouse forebrain, with the highest signal from embryonic day 14 to postnatal day 7, but not in the heart or liver (Supplementary Figure 1). This suggests that the +3 kb enhancer region might be active mainly in neural tissues in late prenatal and early postnatal life.

To further investigate which cell-types the +3 kb region could be active in human *in vivo*, we used data from a recently published human brain cell-type-specific ATAC-seq and ChIP-seq experiments (Nott et al., 2019) (Figure 1B). We found that the +3 kb region shows remarkable neuron specificity, as evident from open chromatin identified using ATAC-seq, and H3K27ac histone mark, which are missing in microglia, oligodendrocytes, and astrocytes. To further elucidate which human tissues and cell types the +3 kb region could be active in, we used the Slidebase tool (Ienasescu et al., 2016) that gathers data of transcription start sites from FANTOM5 consortia and summarizes eRNA transcription levels based on various tissue and cell types. We found that in human the +3 kb region shows the strongest eRNA expression in the brain, spinal cord, and olfactory region (Figure 1C). When grouped by cell type, the strongest expression of +3 kb eRNAs is in hair follicle cells, neurons, and trabecular meshwork cells (Figure 1D).

Collectively, this data suggests that the +3 kb region is an evolutionarily conserved nervous system-specific enhancer that is active mostly in neural tissues and predominantly in neurons but not in other major brain cell types.

### 2. +3 kb enhancer region shows bidirectional transcription in luciferase reporter assay in rat cultured cortical neurons and astrocytes

Based on the enhancer-associated characteristics of the +3 kb enhancer region, we hypothesized that the region could function as an enhancer region for BDNF gene in neural cells. It has been reported that the bidirectional transcription of enhancer RNAs (eRNAs) from the enhancer region is correlated with the expression of nearby genes, indicating that the transcription from an enhancer region is a proxy to enhancer’s activity (Kim et al., 2010). Therefore, we first investigated whether the +3 kb enhancer region shows bidirectional transcription in rat cultured cortical neurons and astrocytes, the two major cell types in the brain, in a heterologous context using reporter assays. We cloned a ~1.4 kb fragment of the +3 kb enhancer and a similarly sized control (+11 kb) intronic sequence lacking enhancer-associated characteristics in either forward or reverse orientation upstream of the firefly luciferase gene (Figure 2A) and performed luciferase reporter assays.

**Figure 2.**
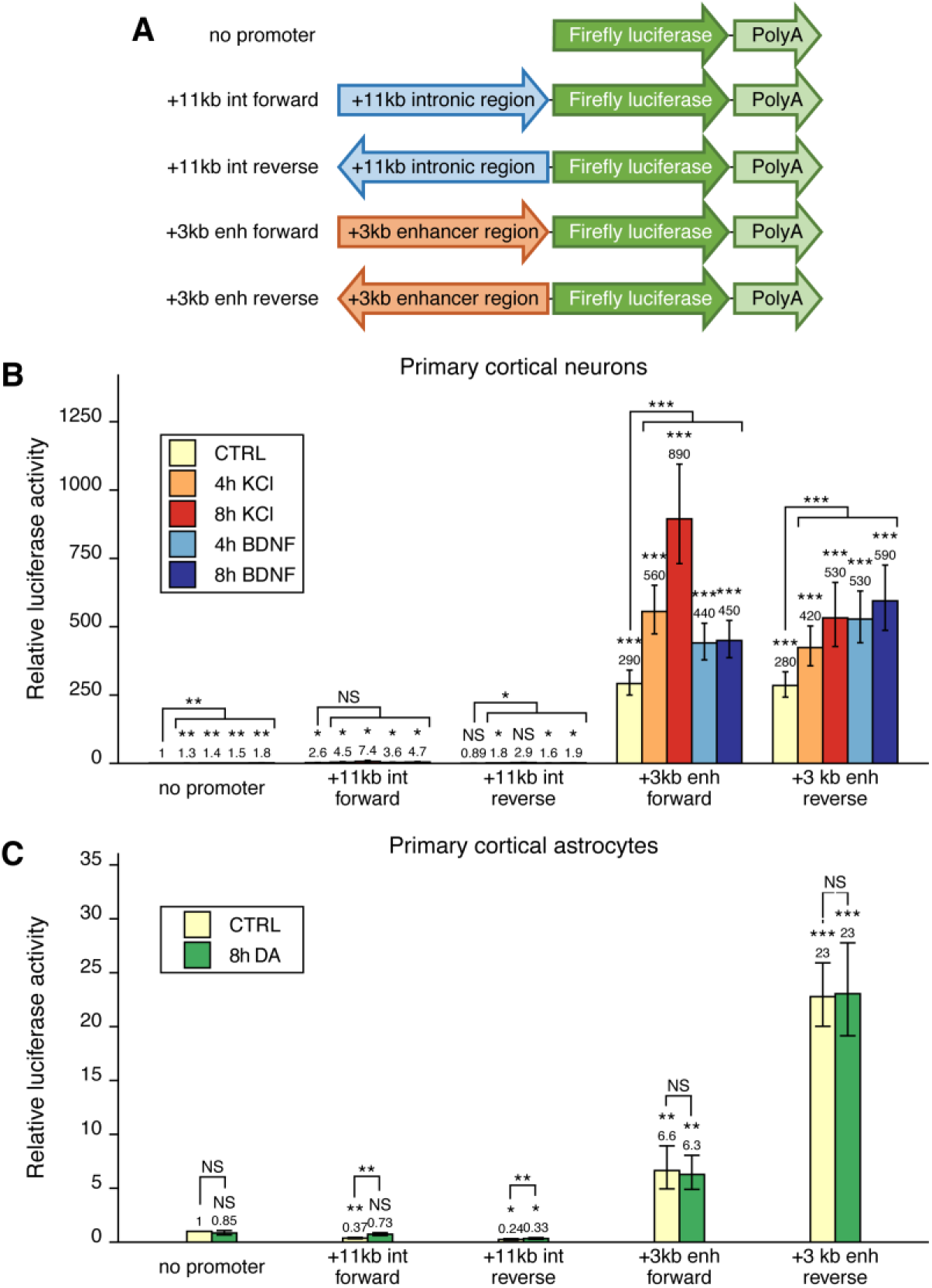
+3 kb enhancer region shows bidirectional transcription in luciferase reporter assay in rat cultured cortical neurons and astrocytes. (A) Reporter constructs used in the luciferase reporter assay where the +3 kb enhancer region and the +11 kb control region were cloned in either forward or reverse orientation (respective to the rat BDNF gene) in front of the luciferase expression cassette. (B, C) Rat cortical neurons (B) or astrocytes (C) were transfected with the indicated reporter constructs at 6 and 13 DIV, respectively. 2 days post transfection, neurons were left untreated (CTRL) or treated with 25 mM KCl (with 5 μM D-APV) or 50 ng/ml BDNF for the indicated time (B); astrocytes were treated with 150 μM dopamine (DA) or respective volume of vehicle (CTRL) for the indicated time (C), after which luciferase activity was measured. Luciferase activity in cells transfected with a vector containing no promoter and treated with vehicle or left untreated was set as 1. The average luciferase activity of independent experiments is depicted above the columns. Error bars indicate SEM (n = 7 (B, +3 kb enhancer constructs and no promoter construct), n = 3 (B, intron constructs), and n = 4 (C) independent experiments). Asterisks above the columns indicate statistical significance relative to luciferase activity in untreated cells transfected with the reporter vector containing no promoter, or between indicated groups. NS – not significant, * p<0.05, ** p<0.01, *** p<0.001 (paired two-tailed t-test).

In rat cortical neurons the +3 kb enhancer region showed very strong transcriptional activity (~300-fold higher compared to the promoterless luciferase reporter vector) that was independent of the orientation of the +3 kb region (Figure 2B). As expected, the +11 kb negative control reporter showed very low luciferase activity in cortical neurons. To determine whether the enhancer region is responsive to different stimuli in neurons and could be involved in stimulus-dependent regulation of the BDNF gene, we used two treatments shown to induce BDNF gene expression – KCl treatment to chronically depolarize the cells and mimic neuronal activity (Ghosh et al., 1994; Pruunsild et al., 2011), and BDNF treatment to activate TrkB signaling and mimic BDNF autoregulation (Esvald et al., 2020; Tuvikene et al., 2016; Yasuda et al., 2007). Our results indicate that the activity of the +3 kb region is upregulated ~2-3-fold in response to both stimuli, suggesting that the region could be a stimulus-dependent enhancer in neurons.

In rat cultured cortical astrocytes, the +3 kb enhancer construct showed modest transcriptional activity (depending on the orientation ~6-23-fold higher compared to the promoterless vector control, Figure 2C). We have previously shown that in cultured cortical astrocytes, BDNF is induced in response to dopamine treatment, and the induction is regulated by an unknown enhancer region within BDNF gene locus (Koppel et al., 2018). However, dopamine treatment had no significant effect on the activity of the +3 kb construct in cultured cortical astrocytes, suggesting that this region is not a dopamine-activated enhancer in astrocytes.

### 3. The +3 kb enhancer region potentiates the activity of BDNF promoters I and IV in luciferase reporter assay in rat cultured cortical neurons

To find out whether the newly identified +3 kb enhancer could control the activity of BDNF promoters in a heterologous context, we cloned the +3 kb region or the +11 kb negative control sequence in forward or reverse orientation (respective to the rat BDNF gene) downstream of BDNF promoter-driven luciferase expression cassette (Figure 3A).

**Figure 3.**
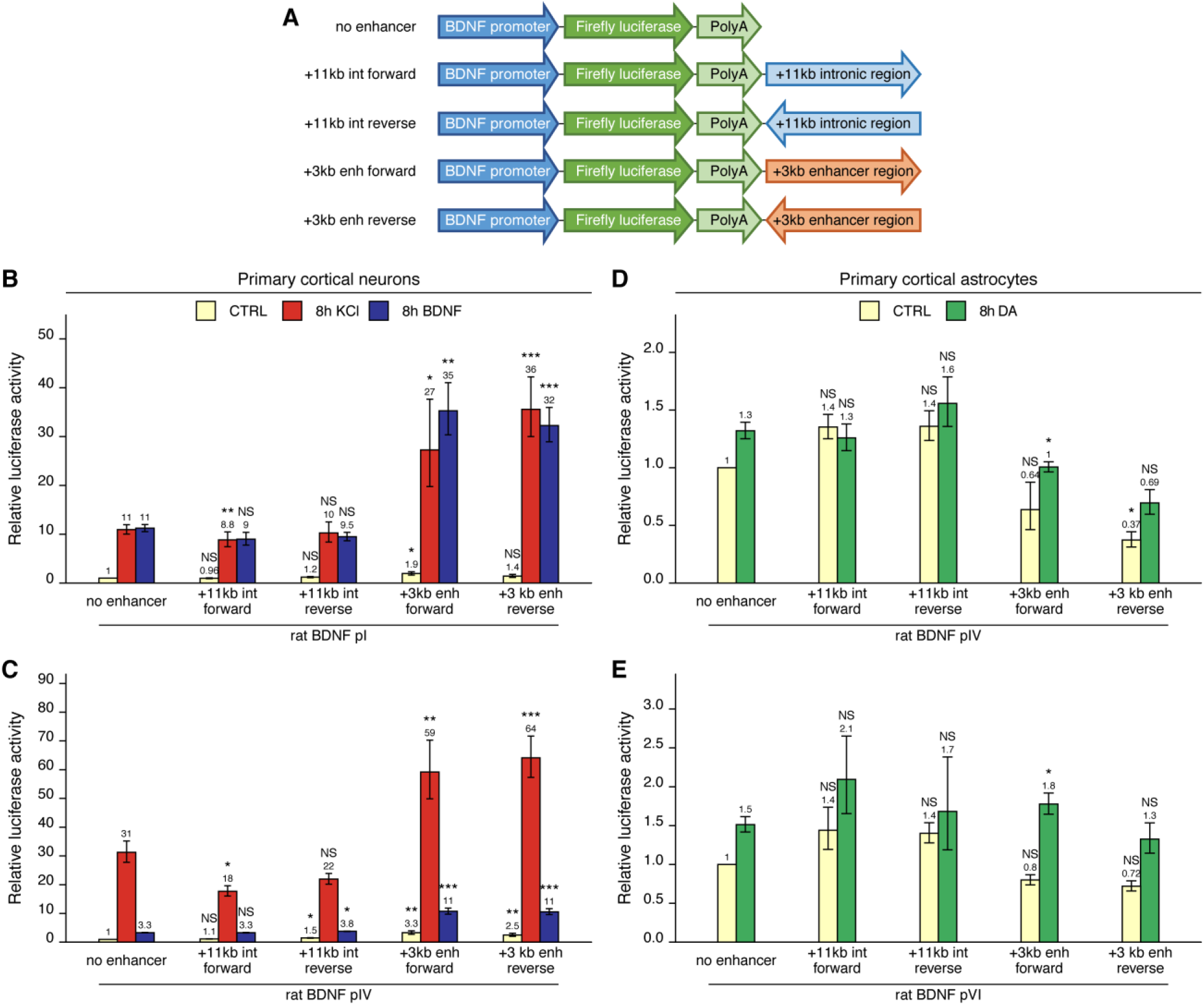
+3 kb enhancer region potentiates the activity of BDNF promoters in luciferase reporter assay in rat cortical neurons but not in astrocytes. (A) A diagram of the used luciferase reporter constructs used in this experiment, with a BDNF promoter in front of the firefly luciferase coding sequence and the +3 kb enhancer or +11 kb intronic region in either forward or reverse orientation (respective to the rat BDNF gene) downstream of the luciferase expression cassette. Rat cortical neurons (B, C) or astrocytes (D, E) were transfected with the indicated reporter constructs at 6 and 13 DIV, respectively. Two days post transfection, neurons were left untreated (CTRL) or treated with 25 mM KCl (with 5 μM D-APV) or 50 ng/ml BDNF for 8 hours (B, C); astrocytes were treated with 150 μM dopamine (DA) or respective volume of vehicle (CTRL) for 8 hours (D, E), followed by luciferase activity assay. Luciferase activity is depicted relative to the luciferase activity in untreated or vehicle-treated (CTRL) cells transfected with respective BDNF promoter construct without an enhancer region. The average luciferase activity of independent experiments is shown above the columns. Error bars represent SEM (n = 6 (B, +3 kb enhancer-containing constructs and no enhancer construct), n = 3 (B, + 11 kb intron constructs), n = 4 (C) and n = 3 (D-E) independent experiments). Statistical significance was calculated compared to the activity of the respective BDNF promoter regions without the enhancer region after the respective treatment. NS – not significant, * p<0.05, ** p<0.01, *** p<0.001 (paired two-tailed t-test).

First, we transfected rat cultured cortical neurons with constructs containing BDNF promoter I and IV, as these promoters are the most widely studied in neurons (West et al., 2014). We treated neurons with either KCl or BDNF, and used luciferase reporter assay to measure the activity of the BDNF promoter region. For BDNF promoter I (Figure 3B), the addition of the +3 kb enhancer region slightly increased the basal activity of the promoter region (~1.5-2-fold). The +3 kb enhancer region also potentiated the KCl and BDNF-induced activity of this promoter ~3-fold in an orientation-independent manner. Similar effects were observed for BDNF promoter IV (Figure 3C), where the addition of the +3 kb enhancer region potentiated the basal activity of the promoter ~3-fold and KCl and BDNF-induced activity levels ~2-fold. The +11 kb intronic region failed to potentiate the activity of BDNF promoters I and IV.

Cortical astrocytes preferentially express BDNF transcripts containing exons IV and VI (Koppel et al., 2018). Therefore, we studied whether the +3 kb enhancer region could promote the activity of BDNF promoters IV and VI in rat cultured cortical astrocytes. The +3 kb enhancer did not significantly increase the activity of BDNF promoters IV and VI in unstimulated astrocytes and had virtually no effect on the response of these promoters to dopamine treatment (Figure 3D, 3E).

Overall, we found that in the heterologous context the +3 kb enhancer region could potentiate transcription from BDNF promoters in cultured cortical neurons, but not in cortical astrocytes. These results imply that the +3 kb enhancer could be important for BDNF gene expression in neurons, but not in astrocytes.

### 4. +3 kb enhancer region is a positive regulator of the first cluster of BDNF transcripts in rat cortical neurons but is in an inactive state in rat cortical astrocytes

To investigate the functionality of the +3 kb region in its endogenous context, we used CRISPR interference (CRISPRi) and activator (CRISPRa) systems. Our system comprised of catalytically inactive Cas9 (dCas9) fused with Krüppel associated box domain (dCas9-KRAB, CRISPRi), or 8 copies of VP16 domain (VP64-dCas9-VP64, CRISPRa) to repress or activate the target region, respectively. dCas9 without effector domains was used to control for potential steric effects (Qi et al., 2013) on BDNF transcription when targeting CRISPR complex inside the BDNF gene. To direct the dCas9 and its variants to the desired location, we used four different gRNAs per region targeting either the +3 kb enhancer region or the +11 kb intronic control region, with all gRNAs targeting the template strand to minimize the potential inhibitory effect of dCas9 binding on transcription elongation (as suggested by Qi et al., 2013). The +11 kb intronic control was used to rule out the possibility of CRISPRi and CRISPRa-induced passive spreading of chromatin modifications within the BDNF gene locus. As a negative control, we used a gRNA not corresponding to any sequence in the rat genome.

We first examined the functionality of the +3 kb enhancer region in cultured cortical neurons. Targeting the +3 kb enhancer or +11 kb intronic region with dCas9 without an effector domain had no major effect on the expression of any of the BDNF transcripts, indicating that targeting CRISPR complex to an intragenic region in BDNF gene does not itself affect BDNF gene expression (Figure 4, left panel). Repressing the +3 kb enhancer region using CRISPRi decreased the basal expression levels of BDNF exon I, IIc, and III-containing transcripts by 2.2, 11, and 2.4-fold, respectively (Figure 4A, 4B, 4C, middle panel). In contrast, no significant effect was seen for basal levels of BDNF exon IV, VI, and IXa-containing transcripts (Figure 4D, 4E, 4F, middle panel). Repressing the +3 kb enhancer region also decreased the KCl and BDNF-induced levels of transcripts starting from the first three 5’ exons ~4-7-fold, but not of other BDNF transcripts (Figure 4A-F, middle panel). These effects correlated with subtle changes in total BDNF expression levels (Figure 4G, middle panel). Targeting CRISPRi to the +11 kb intronic region had no significant effect on any of the BDNF transcripts (Figure 4A-F, middle panel).

**Figure 4.**
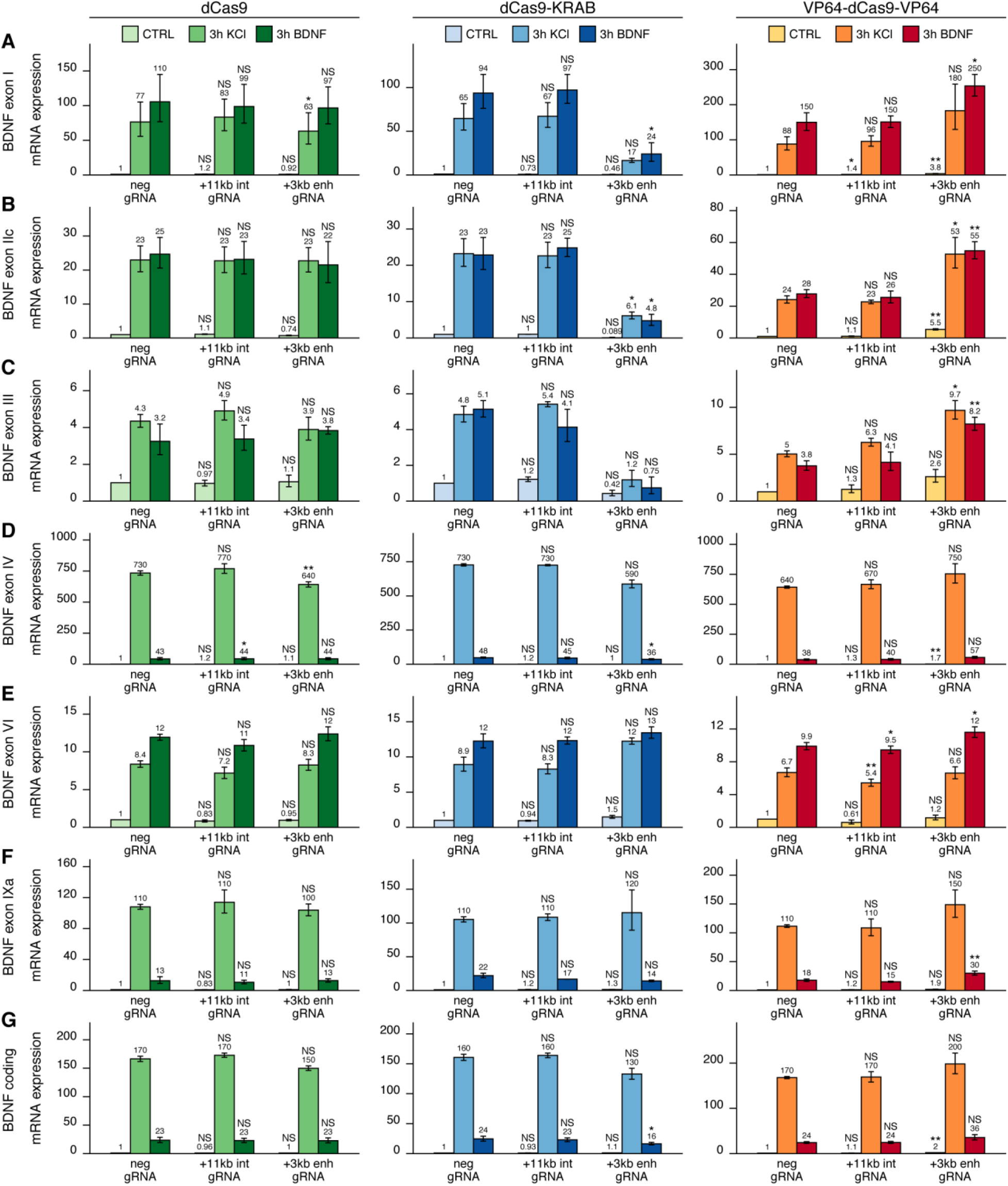
+3 kb enhancer is a positive regulator of BDNF exon I, IIc, and III-containing transcripts in rat cortical neurons. Rat cultured cortical neurons were transduced at 0 DIV with lentiviral particles encoding either catalytically inactive Cas9 (dCas9, left panel, green), dCas9 fused with Krüppel associated box domain (dCas9-KRAB, middle panel, blue) or 8 copies of VP16 domain (VP64-dCas9-VP64, right panel, orange) together with lentiviruses encoding either guide RNA that has no corresponding target sequence in the rat genome (neg gRNA), a mixture of four gRNAs directed to the putative +3 kb BDNF enhancer (+3 kb enh gRNA) or a mixture of four guide RNAs directed to +11 kb intronic region (+11 kb int gRNA). Transduced neurons were left untreated (CTRL) or treated with 50 ng/ml BDNF or 25 mM KCl (with 5 μM D-APV) for 3 hours at 8 DIV. Expression levels of different BDNF transcripts were measured with RT-qPCR. mRNA expression levels are depicted relative to the expression of the respective transcript in untreated (CTRL) neurons transduced with negative guide RNA within each set (dCas9, dCas9-KRAB, or VP64-dCas9-VP64). The average mRNA expression of independent experiments is depicted above the columns. Error bars represent SEM (n = 3 independent experiments). Statistical significance was calculated between the respective mRNA expression levels in respectively treated neurons transduced with neg gRNA within each set (dCas9, Cas9-KRAB, or VP64-dCas9-VP64). NS – not significant, * p<0.05, ** p<0.01, *** p<0.001 (paired two-tailed t-test).

Activating the +3 kb enhancer region with CRISPRa in cultured cortical neurons increased the expression levels of BDNF transcripts of the first cluster (Figure 4A-C, right panel) both in unstimulated neurons (~3-5-fold) and after KCl or BDNF treatment (~2-fold). Slight effect of the activation was also seen for BDNF exon IV and IXa-containing transcripts (Figure 4D, 4F, right panel). Total BDNF basal levels also increased ~2-fold with CRISPR activation of the +3 kb enhancer region, whereas a slightly weaker effect was seen in stimulated neurons. As with CRISPRi, targeting CRISPRa to the +11 kb intronic region had no significant effect on the expression of any of the BDNF transcripts (Figure 4A-F, right panel).

Next, we carried out CRISPRi and CRISPRa experiments in cultured cortical astrocytes. Targeting dCas9 to the BDNF locus did not affect the expression of any of the BDNF transcripts (Figure 5A-F, left panel). Similarly, targeting CRISPRi to the +3 kb enhancer region in astrocytes did not affect the basal expression levels of any BDNF transcript (Figure 5, middle panel). In cells treated with dopamine, repressing +3 kb enhancer region completely abolished the induction of exon IIc-containing transcripts (Figure 5B, middle panel), but did not affect the expression of any of the other transcripts. In contrast, activating the +3 kb region with CRISPRa greatly increased both basal and dopamine-induced levels of all measured BDNF transcripts (Figure 5A-F, right panel). Targeting CRISPRi or CRISPRa to the +11 kb intronic control region did not have a noteworthy effect on any of the measured BDNF transcripts.

**Figure 5.**
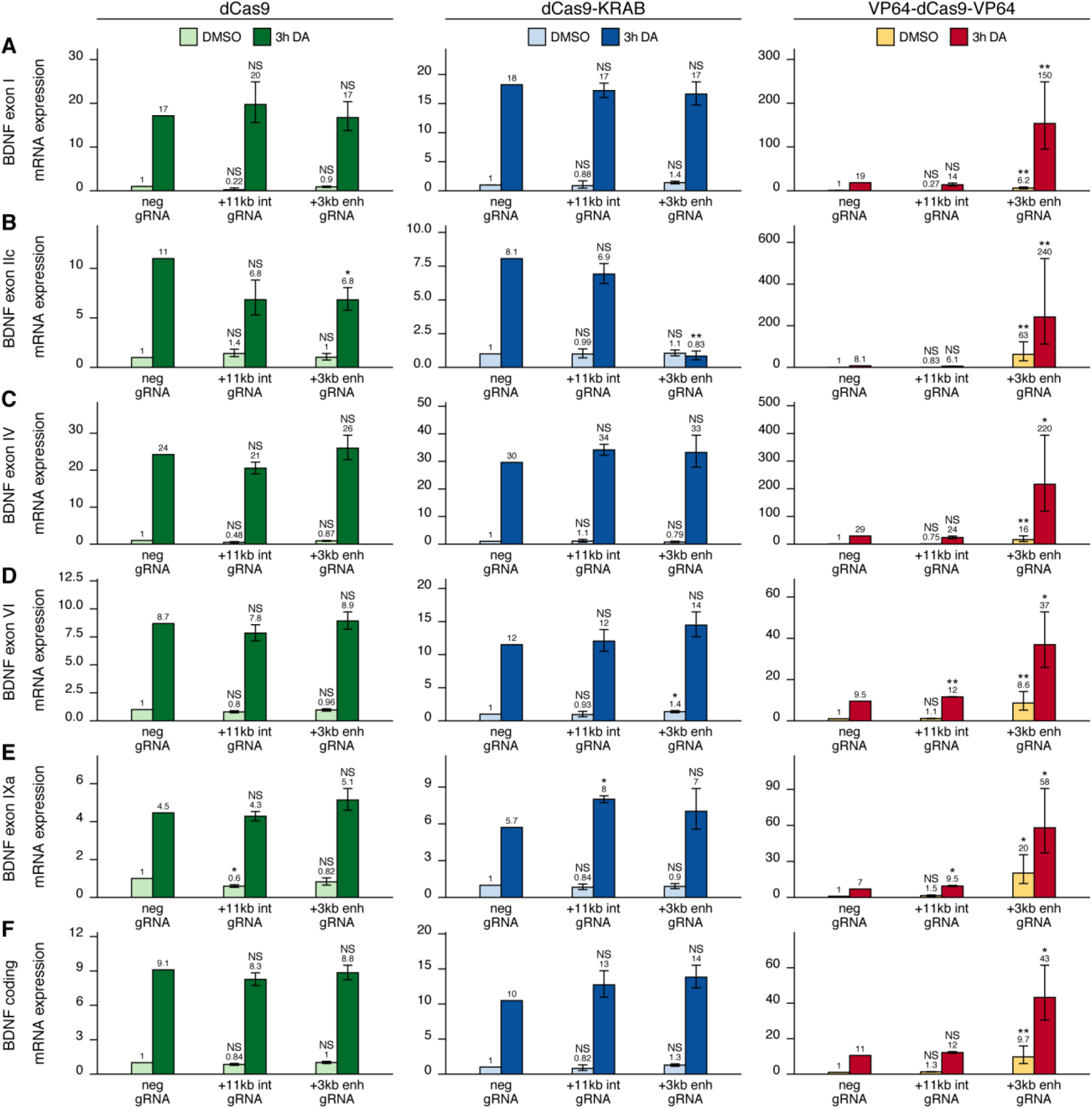
+3 kb enhancer region is mainly inactive in rat cortical astrocytes. Rat cultured cortical astrocytes were transduced at 7 DIV with lentiviral particles encoding either catalytically inactive Cas9 (dCas9, left panel, green), dCas9 fused with Krüppel associated box domain (dCas9-KRAB, middle panel, blue) or 8 copies of VP16 domain (VP64-dCas9-VP64, right panel, orange) together with lentiviruses encoding either guide RNA that has no corresponding target sequence in the rat genome (neg gRNA), a mixture of four gRNAs directed to the putative +3 kb BDNF enhancer (+3 kb enh gRNA) or a mixture of four guide RNAs directed to + 11 kb intronic region (+11 kb int gRNA). Transduced astrocytes were treated with vehicle (CTRL) or with 150 μM dopamine (DA) for 3 hours at 15 DIV. Expression levels of different BDNF transcripts were measured with RT-qPCR. The levels of BDNF exon III-containing transcripts were too low to measure reliably. mRNA expression levels are depicted relative to the expression of the respective transcript in astrocytes treated with vehicle (CTRL) transduced with negative guide RNA within each set (dCas9, dCas9-KRAB, or VP64-dCas9-VP64). The average mRNA expression of independent experiments is depicted above the columns. Error bars represent SEM (n = 3 independent experiments). Statistical significance was calculated between respective mRNA expression levels in respectively treated astrocytes transduced with neg gRNA within each set (dCas9, Cas9-KRAB, or VP64-dCas9-VP64). NS - not significant, * p<0.05, **p<0.01, *** p<0.001 (paired two-tailed t-test).

As bioinformatic analysis showed bidirectional transcription from the +3 kb enhancer (Figure 1) and our luciferase reporter assays also indicated this (Figure 2), we next decided to directly measure eRNAs from the +3 kb enhancer region in our cortical neurons and astrocytes. Since the sense eRNA is transcribed in the same direction as BDNF pre-mRNA, we could only reliably measure eRNAs from the antisense orientation from the +3 kb enhancer region using antisense eRNA-specific cDNA priming followed by qPCR (Supplementary figure 2A). We found that the +3 kb enhancer antisense eRNA was expressed in cultured neurons and the expression level of the eRNA was induced ~3.5- and ~6-fold upon BDNF and KCl treatment, respectively. Furthermore, repressing the +3 kb enhancer region using CRISPRi decreased the expression of the eRNA ~3-fold. However, activating the +3 kb enhancer region using CRISPRa did not change the expression level of the eRNA. When comparing +3 kb enhancer eRNA expression levels in neurons and astrocytes, the astrocytes showed ~6-fold lower eRNA transcription from the +3 kb enhancer region than neurons (Supplementary figure 2B), also indicating that the +3 kb region is in a more active state in our cultured neurons than in astrocytes.

The main findings of the CRISPRi and CRISPRa experiments are summarized in Figure 6. Taken together, our results suggest that the +3 kb enhancer region is an active BDNF enhancer in rat cultured cortical neurons and regulates the basal and stimulus-induced expression of BDNF transcripts of the first cluster of exons (exons I, II and III). In contrast, the +3 kb region is mostly inactive in rat cultured cortical astrocytes.

**Figure 6.**
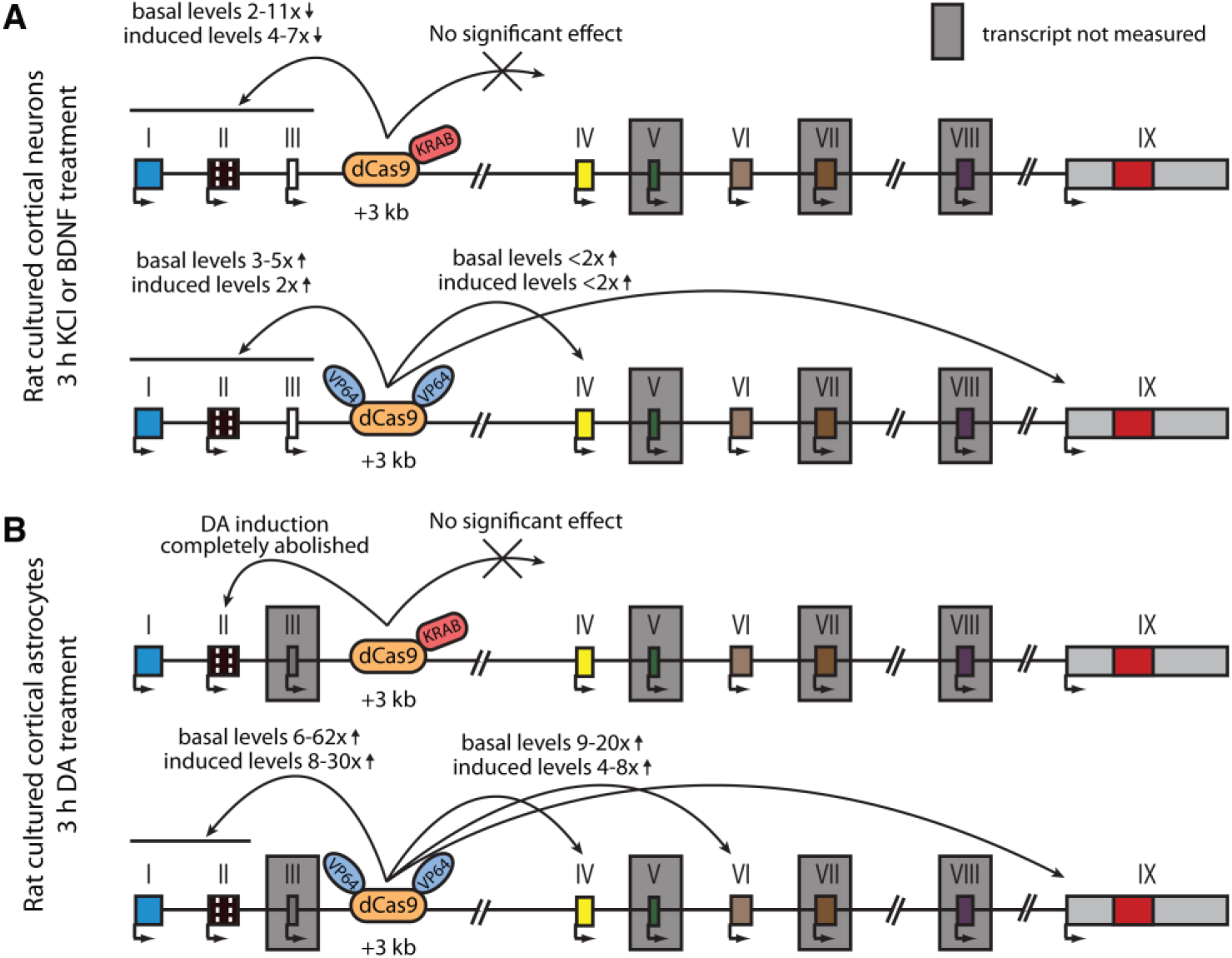
Summary of the CRISPRi and CRISPRa experiments with BDNF +3 kb enhancer region in rat cultured cortical neurons and astrocytes. Graphical representation of the main results shown in Figure 4 (A) and Figure 5 (B). Different BDNF exons are shown with boxes, red box in exon IX indicates BDNF coding region. BDNF transcripts that were not measured or that had too low levels to measure reliably are indicated with a grey box around the respective 5’ exon.

### 5. Deletion of the +3 kb enhancer region in mouse embryonic stem cell-derived neurons decreases the expression of BDNF transcripts starting from the first cluster of exons

To test the regulatory function of the +3 kb enhancer directly and address biological significance of its interspecies conservation, we used CRISPR/Cas9 system to delete the conserved core sequence of the enhancer region in mouse embryonic stem cells engineered to express the pro-neural transcription factor Neurogenin2 from a doxycycline-inducible promoter. We selected single-cell clones containing the desired deletion, differentiated them into excitatory neurons (Ho et al., 2016; Thoma et al., 2012; Zhang et al., 2013) by the addition of doxycyclin, treated the stem cell-derived neurons with BDNF or KCl, and measured the expression of different BDNF transcripts using RT-qPCR.

The deletion of the +3 kb enhancer region strongly decreased both the basal and stimulus-dependent expression levels of BDNF exon I, IIc, and III-containing transcripts (Figure 7A-C). Notably, the effect was more prominent in clones containing homozygous deletion compared to heterozygous clones. We also noted a slight, albeit not statistically significant decrease in the expression of BDNF exon IV and VI-containing transcripts in cells where the +3 kb enhancer region was deleted (Figure 7D-E). This could be attributed to impaired BDNF autoregulatory loop caused by the deficiency of transcripts from the first cluster of exons. It is also possible that the +3 kb enhancer participates in the regulation of the transcripts from the second cluster of exons, but the effect is only very subtle, which would explain why it was not detected in our CRISPRi and CRISPRa experiments in cultured neurons. The levels of BDNF exon IXa were too low to measure reliably (data not shown). The deletion of the +3 kb enhancer region respectively decreased the total levels of BDNF similarly to the first cluster of BDNF transcripts (Figure 7F). These results confirm the essential role of the +3 kb enhancer region in regulating the expression of BDNF exon I, IIc, and III-containing transcripts in rodent neurons.

**Figure 7.**
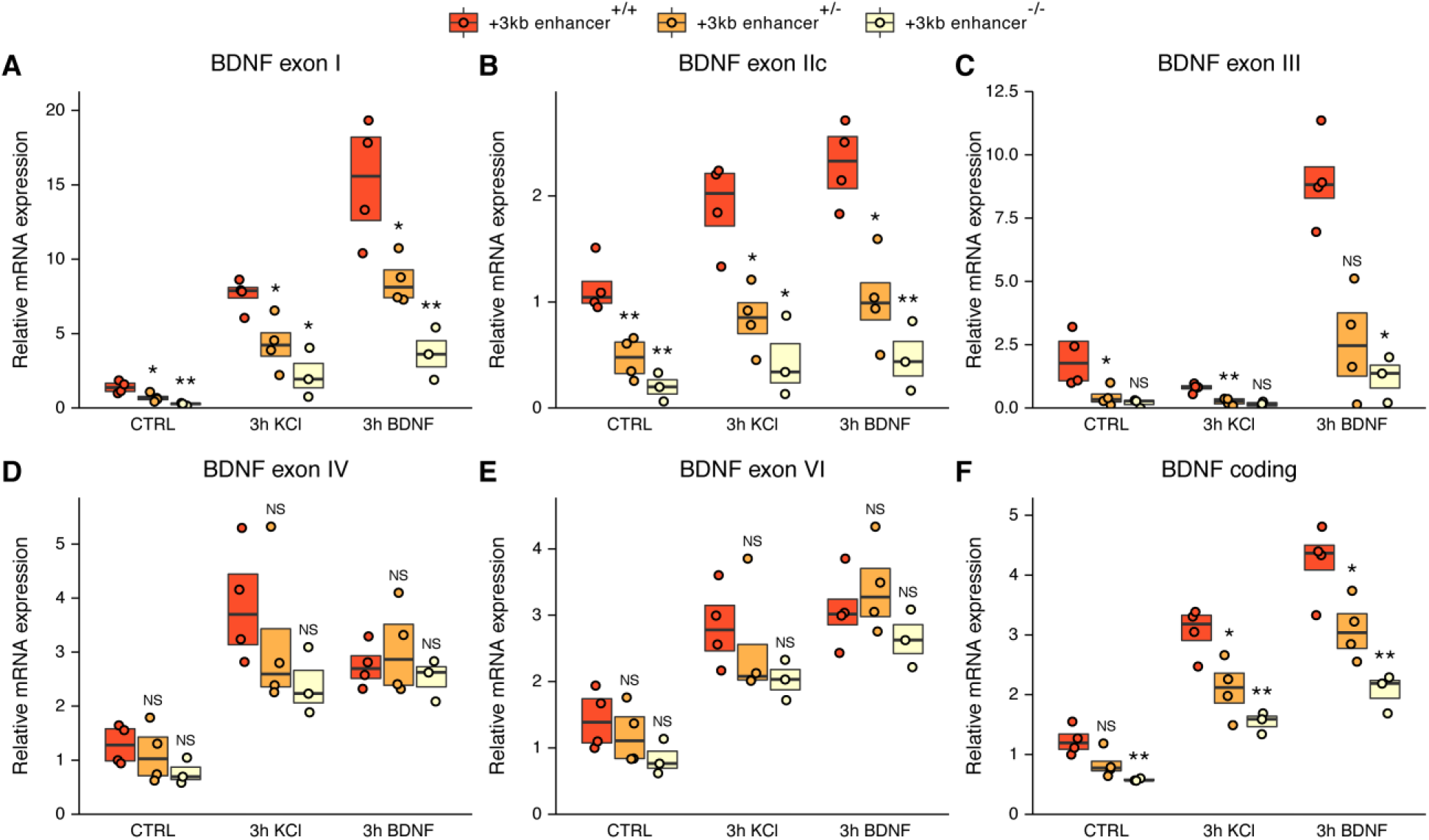
The deletion of the +3 kb enhancer region decreases the expression of BDNF exon I, IIc, and III-containing transcripts in mouse embryonic stem cell (mESC)-derived neurons. CRISPR/Cas9 system was used to generate mESC cell lines with ~300-500 bp deletions of the conserved core region of the +3 kb BDNF enhancer. The obtained clonal cell lines containing intact +3 kb enhancer region (+/+), heterozygous deletion (+/-), or homozygous deletion (-/-) of the +3 kb enhancer region were differentiated into neurons using overexpression of Neurogenin2. After 12 days of differentiation, the cells were treated with vehicle (CTRL), 50 ng/ml BDNF or 25 mM KCl together with 25 μM D-APV for 3 hours. The expression levels of different BDNF transcripts were measured using RT-qPCR. The levels of respective BDNF transcripts measured in the parental cell line (also included as a data point in the +/+ group) was set as 1. All data points (obtained from independent cell clones and parental cell line) are depicted with circles. Box plot shows 25% and 75% quartiles and the horizontal line shows the median value. N = 3-4 independent cell clones for each group. Statistical significance was calculated compared to the expression level of the respective transcript in the +/+ genotype group at respective treatment. NS – not significant, * p < 0.05, ** p < 0.01, *** p < 0.001 (equal variance unpaired t-test).

### 6. The activity of +3 kb enhancer region is regulated by CREB, AP-1 family and E-box-binding transcription factors

To investigate the molecular mechanisms that control the activity of the +3 kb enhancer region, we used *in vitro* DNA pulldown assay with 8 days old rat cortical nuclear lysates coupled with mass-spectrometric analysis to determine the transcription factors that bind to the +3 kb enhancer region (Figure 8A). Collectively, we determined 21 transcription factors that showed specific *in vitro* binding to the +3 kb enhancer region compared to the +11 kb intronic region in two independent experiments (Figure 8B). Of note, we found numerous E-box-binding proteins, including USFs, TCF4 and the pro-neural transcription factors NeuroD2 and NeuroD6, possibly providing the neuron specific activity of the +3 kb enhancer region. We also detected binding of JunD, a member of the AP-1 transcription factor family.

**Figure 8.**
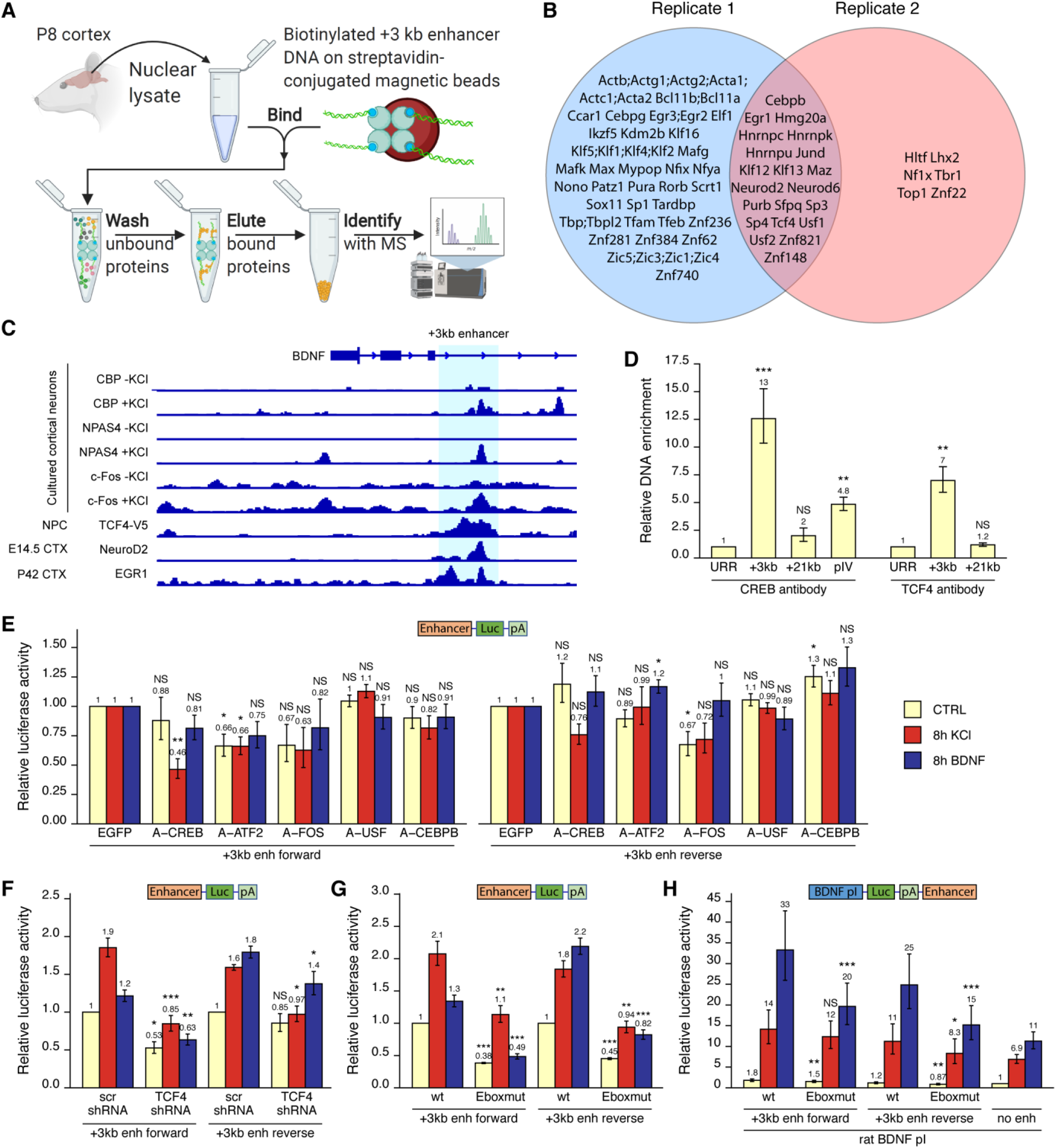
Various transcription factors, including CREB, AP-1 proteins and E-box-binding transcription factors regulate the activity of +3 kb enhancer region. (A) Schematic overview of the in vitro DNA pulldown assay to determine transcription factors binding to the +3 kb enhancer region. The illustration was created with BioRender.com. (B) Transcription factors identified in the in vitro DNA pulldown assay in two biological replicates. Semicolon between protein names indicates uncertainty in the peptide to protein assignment between the proteins separated by the semicolons. (C) Previously published ChIP-seq experiments showing binding of different transcription factors to the +3 kb enhancer region. (D) ChIP-qPCR assay in cultured cortical neurons at 8 DIV with anti-CREB or anti-TCF4 antibody. Enrichment is shown relative to the enrichment of unrelated region (URR) with the respective antibody. +21 kb region (downstream of the BDNF exon VII) was used as a negative control. pIV indicates BDNF promoter IV region. (E-H) Rat cortical neurons were transfected at 5 DIV (F) or 6 DIV (E-H) with reporter constructs where the +3 kb enhancer region was cloned in front of the luciferase coding sequence (E-G, see also Figure 2A) or with reporter constructs where the +3 kb enhancer region was cloned downstream of the BDNF promoter I-controlled firefly luciferase expression cassette (H, see Figure 3A). Schematic representations of the used reporter constructs are shown above the graphs, with Luc designating luciferase coding sequence and pA polyadenylation sequence. At 8 DIV, neurons were left untreated (CTRL) or treated with 25 mM KCl (with 5 μM D-APV) or 50 ng/ml BDNF for the indicated time, after which luciferase activity was measured. Luciferase activity is depicted relative to the luciferase activity in respectively treated cells transfected with EGFP and the respective +3 kb enhancer construct (E), relative to luciferase activity in untreated cells co-transfected with control shRNA (scr) and the respective +3 kb enhancer construct (F), relative to luciferase activity in untreated cells transfected with the respective wild-type (wt) +3 kb enhancer construct (G), or relative to luciferase activity in untreated cells transfected with rat BDNF promoter I construct containing no enhancer region (H). Eboxmut indicates mutation of a putative E-box element in the +3 kb enhancer region. Numbers above the columns indicate average, error bars represent SEM (n = 4 (D, TCF4 antibody), n = 3 (D, CREB antibody), n = 5-6 (E), n = 4-5 (F), n=4 (G), n = 7 (H) independent experiments). Statistical significance was calculated compared to the ChIP enrichment of DNA at the URR region using respective antibody (D), compared to the luciferase activity in respectively treated cells transfected with respective +3 kb enhancer construct and EGFP (E), scr shRNA (F), or respective wt +3 kb enhancer construct (G, H). NS – not significant, * p<0.05, ** p<0.01, *** p<0.001 (paired two-tailed t-test).

Next, we used various ChIP-seq experiments in different human cell lines from the ENCODE project and determined numerous transcription factors that bind to the +3 kb enhancer region, including CREB, CEBPB, EGR1 and JunD (Supplementary figure 3). We further used publicly available ChIP-seq data (see Materials and methods section for references) to visualize the binding of different transcription factors to the +3 kb enhancer region in mouse neural cells and tissues (Figure 8C). This data shows neuronal activity-dependent binding of NPAS4, c-Fos, and coactivator CBP to the enhancer region. In agreement with our *in vitro* pulldown results, ChIP-seq analysis also revealed binding of EGR1, NeuroD2 and TCF4 to the +3 kb enhancer region in the endogenous chromatin context.

Considering the CREB binding in ENCODE data (Suplementary figure 2) and CBP binding in cultured cortical neurons (Figure 8C), we first decided to investigate whether CREB binds BDNF +3 kb enhancer region in our rat cortical neurons. We performed ChIP-qPCR and determined that in cultured cortical neurons CREB binds to the +3 kb enhancer region, whereas we found no significant CREB binding to the +21 kb negative control region, located directly downstream of BDNF exon VII (Figure 8D). Of note, the binding of CREB to the +3 kb enhancer region was ~2.6-times stronger than binding to BDNF promoter IV, which contains the well-described CRE element (Hong et al., 2008; Tao et al., 1998). Next, we focused on the various E-box-binding proteins, as many E-box-binding proteins are proneural and could therefore confer the neural specificity of the +3 kb enhancer region. As transcription factors from the NeuroD family need dimerization partner from the class I helix-loop-helix proteins, e.g. TCF4, to bind DNA (Massari and Murre, 2000; Ravanpay and Olson, 2008), we verified binding of TCF4 to the +3 kb region in our cultured neurons using TCF4 ChIP-qPCR (Figure 8D).

To determine functionally important transcription factors that regulate BDNF +3 kb enhancer region, we first screened a panel of dominant-negative transcription factors in luciferase reporter assay where the expression of luciferase was under control of the +3 kb enhancer region (Figure 8E). In agreement with the in *vitro* pulldown assay, ChIP-seq and ChIP-qPCR results, we found the strongest inhibitory effect using dominant negative versions of CREB (named A-CREB), ATF2 (named A-ATF2) and AP-1 family (named A-FOS). However, the effect of different dominant negative proteins was slightly lower when +3 kb enhancer region was in the reverse orientation. Our data suggests the role of CREB, AP-1 family proteins and ATF2 in regulating the neuronal activity-dependent activation of the +3 kb enhancer region, whereas we found no notable evidence of USF family transcription factors and CEBPB regulating the activity of +3 kb enhancer region.

Finally, we elucidated the role of E-box binding proteins in the regulation of +3 kb enhancer region. Using luciferase reporter assays, we found that silencing TCF4 expression with TCF4 shRNA-expressing plasmid decreased the activity of the +3 kb enhancer region in both unstimulated and KCl and BDNF-stimulated neurons. However, the effects were slightly smaller when the enhancer region was in the reverse orientation (Figure 8F). Based on the TCF4 and NeuroD2 ChIP-seq data (Figure 8C), we identified a putative E-box binding sequence in the +3 kb enhancer region (CAGATG). To determine the relevance of this E-box element, we generated +3 kb enhancer-containing reporter constructs where this E-box motif was mutated (CAGAAC). We determined that this motif participates in regulating both the basal activity and BDNF and KCl-induced activity of the enhancer region (Figure 8G). Importantly, mutating the E-box decreased the ability of the +3 kb enhancer region to potentiate transcription from BDNF promoter I in reporter assays (Figure 8H).

Collectively, we have identified numerous transcription factors that potentially regulate the activity of +3 kb enhancer region, and further discovered a functional E-box element in the enhancer, possibly conferring neuron-specific activity of the +3 kb enhancer region.

## DISCUSSION

BDNF promoters I, II, and III are located within a relatively compact (~2 kb) region in the genome, making it possible that their activity is controlled by a common mechanism. A similar spatial clustering of BDNF exons seems to be conserved in vertebrates, with a similar genomic organization observed in frog (Kidane et al., 2009), chicken (Yu et al., 2009), zebrafish (Heinrich and Pagtakhan, 2004), rodents and human (Aid et al., 2007; Pruunsild et al., 2007). It has previously been suggested that BDNF promoters I and II could be co-regulated as one functional unit (Hara et al., 2009; Timmusk et al., 1999; West et al., 2014). Here, we show that the promoters of BDNF exons I, II, and III are co-regulated as a neuron-specific unit through a conserved enhancer region located downstream of exon III.

We have previously reported that exon I-containing BDNF transcripts contain in-frame alternative translation start codon that is used more efficiently for translation initiation than the canonical start codon in the exon IX (Koppel et al., 2015). As the BDNF exon I-containing transcripts are highly inducible in response to different stimuli, they could make a substantial contribution to the overall production of BDNF protein in neurons, despite the low basal expression levels of this transcript. Remarkably, the BDNF transcripts from the first cluster of exons have been shown to regulate important aspects of behavior. In female mice, BDNF exon I-containing transcripts are important for proper sexual and maternal behavior (Maynard et al., 2018b), whereas in male mice the BDNF exon I and exon II-containing transcripts regulate serotonin signaling and control aggressive behavior (Maynard et al., 2016b). Furthermore, it has been shown that BDNF exon I-containing transcripts in the hypothalamus participate in energy metabolism and thermoregulation (You et al., 2020). The +3 kb enhancer identified in our work might therefore be an important regulator of BDNF gene expression in the formation of the neural circuits regulating both social behavior and energy metabolism. Further work will address this possibility experimentally.

The data from Nord et al. (2013) indicates that the highest H3K27ac modification, a hallmark of active regulatory region, at the +3 kb enhancer region in development occurs a week before and a week after birth in mice – coinciding with the period of late neurogenesis, neuronal migration, synaptogenesis, and maturation of neurons (Reemst et al., 2016). It appears that the +3 kb enhancer region is mostly active in early life and participates in the development of the central nervous system via regulating BDNF expression. However, it is also possible that the decline in H3K27ac mark in murine brain tissue during postnatal development is due to the increased amount of non-neuronal cells in the brain compared to neurons. Although the activity of the +3 kb enhancer seems to decrease with age, it is plausible that it remains active also in later postnatal life and upregulates BDNF expression, thereby regulating synaptic plasticity in the adult organism.

Based on the induction of eRNA expression from the +3 kb enhancer region upon depolarization and BDNF-TrkB signaling in both luciferase reporter assays and in the endogenous context, and binding of various activity-dependent transcription factors to the +3 kb region, our data indicates that in addition to conferring neuron-specificity, the +3 kb enhancer region also participates in BDNF-TrkB signaling and neuronal activity-induced expression of the first cluster of BDNF transcripts. Furthermore, repressing or activating the +3 kb enhancer region with CRISPRi or CRISPRa also affected the stimulus-induced levels of these transcripts. Notably, the part of the BDNF gene containing the +3 kb enhancer has previously been implicated in the Reelin-mediated induction of BDNF expression (Telese et al., 2015), indicating that the +3 kb enhancer could respond to other stimuli in addition to membrane depolarization and TrkB signaling.

We also investigated the possibility that the +3 kb enhancer contributes to the catecholamine-induced expression of BDNF transcripts in rat cultured cortical astrocytes (Koppel et al., 2018) and noted that even though the activation of the +3 kb enhancer increased the basal and stimulus-induced expression of all BDNF transcripts, repression of the +3 kb enhancer had almost no effect on BDNF expression. Furthermore, the transcriptional activity of the +3 kb enhancer was not induced by dopamine-treatment in luciferase reporter assay, further indicating that the +3 kb region is not the enhancer responsible for catecholamine-dependent induction of BDNF expression. Interestingly, the dopamine-dependent induction of BDNF exon II-containing transcripts was abolished when the +3 kb enhancer was repressed using CRISPRi, suggesting that the +3 kb enhancer region might control the activity of stimulus-specific expression of BDNF promoter II in astrocytes. Since the activity of BDNF promoter II is regulated by neuron-restrictive silencer factor (NRSF) (Timmusk et al., 1999), it is possible that the drastic decrease in the dopamine-dependent induction of BDNF exon II-containing transcripts was due to the cooperative effect between NRSF and the +3 kb enhancer region. Further investigation is needed to determine whether this hypothesis is true and whether such cooperation between the +3 kb enhancer region and NRSF binding to BDNF exon II also happens in neurons. Although we have not tested this directly, our data does not support the notion that the +3 kb region is an active repressor in non-neuronal cells, e.g. astrocytes. Instead, it seems that the +3 kb enhancer is a positive regulator of BDNF gene operating specifically in neurons. We conclude that the +3 kb enhancer region is largely inactive in rat cultured cortical astrocytes and it is distinct from the distal *cis*-regulatory region controlling the catecholamine-induced activities of BDNF promoters IV and VI.

Our results indicate that the +3 kb enhancer can receive regulatory inputs from various basic helix-loop-helix transcription factors, including TCF4 and its pro-neural heterodimerization partners NeuroD2 and NeuroD6. Single-cell RNA-seq analysis in the mouse cortex and hippocampus has indicated that NeuroD transcription factors are expressed mainly in excitatory neurons, similar to BDNF (Tasic et al., 2016). It has been reported that NeuroD2 preferentially binds to E-boxes CAGCTG or CAGATG (Fong et al., 2012), which is in agreement with the functional E-box CAGATG sequence found in the +3 kb enhancer. Furthermore, it has been previously shown that NeuroD2 knock-out animals exhibit decreased BDNF levels in the cerebellum (Olson et al., 2001). However, Olson et al found no change in BDNF levels in the cerebral cortex of these knock-out animals. It is possible that different NeuroD family transcription factors regulate BDNF expression in different brain areas and developmental stages, or that a compensatory mechanism between NeuroD2 and NeuroD6, both binding the +3 kb enhancer region in our *in vitro* DNA pulldown assay, exists in cortical neurons of NeuroD2 knock-out animals. It has been well-described that NeuroD transcription factors regulate neuronal differentiation (Massari and Murre, 2000), axonogenesis (Bormuth et al., 2013), neuronal migration (Guzelsoy et al., 2019), and proper synapse formation (Ince-Dunn et al., 2006; Wilke et al., 2012). As BDNF also has a role in the aforementioned processes (Park and Poo, 2013), it is plausible that at least some of the effects carried out by NeuroD family result from increasing BDNF expression. Further work is needed to clarify the exact role of TCF4 and NeuroD transcription factors in BDNF expression.

In conclusion, we have identified a novel intronic enhancer region governing the expression of neuron-specific BDNF transcripts starting from the first cluster of exons – exons I, II, and III – in mammals. Exciting questions for further work are whether the +3 kb enhancer region is active in all neurons or in specific neuronal subtypes, and whether the activity of this enhancer element underlies *in vivo* contributions of BDNF to brain development and function.

## MATERIALS AND METHODS

### 1. Cultures of rat primary cortical neurons

All animal procedures were performed in compliance with the local ethics committee. Cultures of cortical neurons from embryonic day (E) 21 Sprague Dawley rat embryos of both sexes were prepared as described previously (Esvald et al., 2020). The cells were grown in Neurobasal A (NBA) medium (Gibco) containing 1× B27 supplement (Gibco), 1 mM L-glutamine (Gibco), 100 U/ml penicillin and 0.1 mg/ml streptomycin (Gibco) or 100 μg/ml Primocin (Invivogen) instead of penicillin/streptomycin at 37 °C in 5% CO_2_ environment. At 2 days *in vitro* (DIV), half of the medium was replaced with fresh supplemented NBA, or the whole medium was replaced for cells transduced with lentiviruses. To inhibit the proliferation of non-neuronal cells a mitotic inhibitor 5-fluoro-2’-deoxyuridine (FDU, final concentration 10 μM, Sigma-Aldrich) was added with the change of the medium.

### 2. Cultures of rat primary cortical astrocytes

Cultures of cortical astrocytes were prepared from E21 Sprague Dawley rat embryos of both sexes as described previously (Koppel et al., 2018). The cells were grown in 75 cm^2^ tissue culture flasks in Dulbecco’s Modified Eagle Medium (DMEM with high glucose, PAN Biotech) supplemented with 10% fetal bovine serum (PAN Biotech) and 100 U/ml penicillin and 0.1 mg/ml streptomycin (Gibco) at 37 °C in 5% CO_2_ environment. At 1 DIV, the medium was replaced with fresh growth medium to remove loose tissue clumps. At 6 DIV, the flasks were placed into a temperature-controlled shaker Certomat^®^ BS-1 (Sartorius Group) for 17-20 hours and shaken at 180 rpm at 37 °C to detach non-astroglial cells from the flask. After overnight shaking, the medium was removed along with unattached non-astrocytic cells, and astrocytes were washed three times with 1× PBS. Astrocytes were detached from the flask with trypsin-EDTA solution (0.25% Trypsin-EDTA (1×), Gibco) diluted 4 times with 1× PBS at 37 °C for 3-5 minutes. Trypsinized astrocytes were resuspended in supplemented DMEM and centrifuged at 200 × g for 6 minutes. The supernatant was removed, astrocytes were resuspended in supplemented DMEM and seeded on cell culture plates previously coated with 0.2 mg/ml poly-L-lysine (Sigma-Aldrich) in Milli-Q. At 9 DIV, the whole medium was replaced with fresh supplemented DMEM.

### 3. Drug treatments

At 7 DIV, cultured neurons were pre-treated with 1 μM tetrodotoxin (Tocris) until the end of the experiment to inhibit spontaneous neuronal activity. At 8 DIV, neurons were treated with 50 ng/ml human recombinant BDNF (Peprotech) or with a mixture of 25 mM KCl and 5 μM NMDA receptor antagonist D-2-amino-5-phosphopentanoic acid (D-APV, Cayman Chemical Company) to study BDNF autoregulation or neuronal activity-dependent expression of the BDNF gene, respectively.

Cultured cortical astrocytes were treated at 15 DIV with 150 μM dopamine (Tocris) to study the regulation of the BDNF gene by catecholamines or 0.15% DMSO (Sigma) as a vehicle control in fresh serum-free and antibiotics-free DMEM (DMEM with high glucose, PAN Biotech).

### 4. Transfection of cultured cells and luciferase reporter assay

Rat +3 kb enhancer (chr3:100771267-100772697, rn6 genome assembly) or +11 kb intron (chr3:100778398-100779836, rn6 genome assembly) regions were amplified from rat BDNF BAC construct (Koppel et al., 2018) using Phusion Hot Start II DNA Polymerase (Thermo Fisher Scientific) and cloned into pGL4.15 vector (Promega) in front of the Firefly luciferase coding sequence. To generate reporter constructs containing both BDNF promoter and enhancer region, the hygromycin expression cassette downstream of Firefly luciferase expression cassette in pGL4.15 vector was replaced with a new multiple cloning site, into which the +3 kb enhancer or +11 kb intron regions were cloned in either forward or reverse orientation (respective to the rat BDNF gene). The BDNF promoter regions were obtained from rat BDNF promoter constructs (Esvald et al., 2020) and cloned in front of the Firefly luciferase coding sequence. Plasmids encoding control and TCF4 shRNA have been published previously (Sepp et al., 2017). Coding regions of different dominant negative transcription factors were subcloned from AAV plasmids (Esvald et al., 2020) into pRRL vector backbone under the control of human PGK promoter.

For transfection and luciferase reporter assays, rat cortical neurons or astrocytes were grown on 48-well cell culture plates. Transfections were carried out in duplicate wells.

Cultured cortical neurons were transfected as described previously (Jaanson et al., 2019) with minor modifications. Transfection was carried out in unsupplemented NBA using 500 ng of the luciferase reporter construct and 20 ng of a normalizer plasmid pGL4.83-mPGK-hRLuc at 5-6 DIV using Lipofectamine 2000 (Thermo Scientific) with DNA to Lipofectamine ratio of 1:2. Transfection was terminated by replacing the medium with conditioned medium, which was collected from the cells before transfection.

Cultured cortical astrocytes were transfected as described previously (Koppel et al., 2018) using 190 ng of luciferase reporter construct and 10 ng of normalizer plasmid pGL4.83-SRα-hRLuc at 13 DIV using Lipofectamine 2000 (Thermo Scientific) with DNA to Lipofectamine ratio of 1:3.

The cells were lysed with Passive Lysis Buffer (Promega) and luciferase signals were measured with Dual-Glo^®^ Luciferase assay kit (Promega) using GENios pro plate reader (Tecan). Background-corrected Firefly luciferase signals were normalized to background-corrected Renilla luciferase signals and the averages of duplicate wells were calculated. Data were log-transformed for statistical analysis, mean and standard error of the mean (SEM) were calculated, and data were back-transformed for graphical representation.

### 5. CRISPR interference and activator systems, RT-qPCR

pLV-hUbC-dCas9-KRAB-T2A-GFP plasmid used for CRISPR interference has been described previously (Esvald et al., 2020) and pLV-hUbC-VP64-dCas9-VP64-T2A-GFP plasmid used for CRISPR activation was obtained from Addgene (plasmid #59791). Lentiviral particles were produced as described previously (Koppel et al., 2018). Relative viral titers were estimated from provirus incorporation rate measured by qPCR and an equal amounts of functional viral particles were used for transduction in the following experiments.

Rat cortical neurons were transduced at 0 DIV, whereas cortical astrocytes were transduced after sub-culturing at 7 DIV. After treatments at 8 DIV for neurons or at 14 DIV for astrocytes, the cells were lysed and RNA was extracted with RNeasy Mini Kit (Qiagen) using on-column DNA removal with RNase-Free DNase Set (Qiagen). RNA concentration was measured with BioSpec-nano spectrophotometer (Shimadzu Biotech). cDNA was synthesized from equal amounts of RNA with Superscript^®^ III or Superscript^®^ IV reverse transcriptase (Invitrogen) using oligo(dT)20 or a mixture of oligo(dT)20 and random hexamer primer (ratio 1:1, Microsynth). To measure +3 kb enhancer eRNAs, cDNA was synthesized using a mixture of antisense-eRNA specific primer and HPRT1 primer (1:1 ratio). The primers used for cDNA synthesis are listed in Supplementary Table 1.

All qPCR reactions were performed in 10 μl volume in triplicates with 1× HOT FIREpol EvaGreen qPCR Mix Plus (Solis Biodyne) and primers listed in Supplementary Table 1 on LightCycler^®^ 480 PCR instrument II (Roche). Gene expression levels were normalized to HPRT1 mRNA levels in neurons and Cyclophilin B mRNA levels in astrocytes. Data were log-transformed and autoscaled (as described in Vandesompele et al., 2002) for statistical analysis, mean and SEM were calculated, and data were back-transformed for graphical representation.

### 6. Mouse embryonic stem cells

A2Lox mouse embryonic stem cells (mESCs) containing doxycyclin-inducible Neurogenin2 transgene (Zhuravskaya and Makeyev, in preparation) were grown in 2i media as described in (Iacovino et al., 2011; Kainov and Makeyev, 2020). To delete the +3 kb enhancer region, 3 + 3 gRNAs targeting either side of the +3 kb enhancer core region (targeting sequences listed in Supplementary Table 2) were cloned into pX330 vector (Addgene plasmid #42230). mESCs were co-transfected with a mixture of all 6 CRISPR plasmids and a plasmid containing a blasticidin expression cassette for selection. One day post transfection, 8 μg/ml blasticidin (Sigma) was added to the media for 3 days after which selection was ended, and cells were grown for an additional 11 days. Finally, single colonies were picked and passaged. The deletion of the +3 kb enhancer region was assessed from genomic DNA with PCR using primers flanking the desired deletion area. To rule out larger genomic deletions, qPCR-based copy number analysis was carried out with primers targeting the desired deletion area, and either side of the +3 kb region outside of the desired deletion area. All primers are listed in Supplementary Table 1. Cell clones containing no deletion, heterozygous or homozygous deletion of the core conserved enhancer region together with intact flanking regions were used for subsequent analysis.

Selected mESCs were differentiated into neurons as follows. Cells were plated on Matrigel (Gibco) coated 12-well plates at a density of ~25 000 cells/well in N2B27 media (1:1 DMEM F12-HAM and Neurobasal mixture, 1x N2, 1x B27 with retinoic acid, 1x penicillin-streptomycin, 1 μg/ml laminin, 20 μg/ml insulin, 50 μM L-glutamine) supplemented with 0.1M β-mercaptoethanol (Sigma) and 2 μg/ml doxycycline (Sigma). After 2 days the whole media was changed to N2B27 media containing 200 μM ascorbic acid (Sigma) and 1 μg/ml doxycycline. Next, half of the media was replaced every 2 days with new N2B27 media containing 200 μM ascorbic acid but no doxycycline. On the 12^th^ day of differentiation, cells were treated with Milli-Q, BDNF (Peprotech), or KCl for 3 hours. All treatments were added together with 25 μM D-APV (Alfa Aesar). After treatment, the cells were lysed and RNA was extracted using EZ-10 DNAaway RNA Mini-Prep Kit (Bio Basic inc). cDNA was synthesized using Superscript IV (Thermo Fischer) and qPCR was performed with HOT FIREPol EvaGreen qPCR Mix Plus (Solis Biodyne) or qPCRBIO SyGreen Mix Lo-ROX (PCR Biosystems Ltd) on LightCycler 96 (Roche). The levels of CNOT4 mRNA expression were used for normalization. Used primers are listed in Supplementary Table 1.

### 7. In vitro DNA pulldown mass-spectrometry

855 bp region of the +3 kb enhancer and +11 kb intronic region were amplified with PCR using HotFirePol polymerase (Solis Biodyne) and primers listed in Supplementary Table 1, with the reverse primers having a 5’ biotin modification (Microsynth). PCR products were purified using DNA Clean & Concentrator™-100 kit (Zymo Research) using a 1:5 ratio of PCR solution and DNA binding buffer. The concentration of the DNA was determined with Nanodrop 2000 spectrophotometer (Thermo Scientific).

The preparation of nuclear lysates was performed as follows. Cortices from 8 days old Sprague Dawley rat pups of both sexes were dissected and snap-frozen in liquid nitrogen. Nuclear lysates were prepared with high salt extraction as in (Wu, 2006) and (Lahiri and Ge, 2000) with minor modifications. Briefly, cortices were weighed and transferred to pre-cooled Dounce tissue grinder (Wheaton). 2 ml of ice-cold cytoplasmic lysis buffer (10 mM Hepes, pH 7.9 (adjusted with NaOH), 10 mM KCl, 1.5 mM MgCl_2_, 0.5% NP-40, 300 mM sucrose, 1x cOmplete™ Protease Inhibitor Cocktail (Roche), and phosphatase inhibitors as follows: 5 mM NaF (Fisher Chemical), 1 mM beta-glycerophosphate (Acros Organics), 1 mM Na_3_VO_4_ (ChemCruz, sc-3540A) and 1 mM Na_4_P_2_O_7_ (Fisher Chemical)) was added and tissue was homogenized 10 times with tight pestle. Next, the lysate was transferred to a 15 ml tube and cytoplasmic lysis buffer was added to a total volume of 1 ml per 0.1 g of tissue. The lysate was incubated on ice for 10 minutes with occasional inverting. Next, the lysate was transferred to a 100 μm nylon cell strainer (VWR, ref nr 732-2759) to remove tissue debris and the flow-through was centrifuged at 2600 × g at 4 °C for 1 min to pellet nuclei. The supernatant (cytoplasmic fraction) was discarded and the nuclear pellet was resuspended in 1 ml per 1 g of tissue ice-cold nuclear lysis buffer (20 mM Hepes, pH 7.9, 420 mM NaCl, 1.5 mM MgCl_2_, 0.1 mM EDTA, 2.5% Glycerol, 1x cOmplete™ Protease Inhibitor Cocktail (Roche), and phosphatase inhibitors) and transferred to a new Eppendorf tube. To extract nuclear proteins, the pellet was rotated at 4 °C for 30 minutes and finally centrifuged at 11 000 × g at 4 °C for 10 min. The supernatant was collected as nuclear fraction and protein concentration was measured with BCA Protein Assay Kit (Pierce).

*In vitro* DNA pulldown was performed as follows. Two biological replicates were performed using nuclear lysates of cortices from pups of different litters. Pierce™ Streptavidin Magnetic Beads (50 μl per pulldown reaction) were washed 2 times with 1× binding buffer (BB, 5 mM Tris-HCl, pH 7.5, 0.5 mM EDTA, 1 M NaCl, 0.05% Tween-20), resuspended in 2× BB, and an equal volume of 50 pmol biotinylated DNA (in 10 mM Tris-HCl, pH 8.5, 0.1 mM EDTA) was added and incubated at room temperature for 30 min with rotation. To remove the unbound probe, the beads were washed 3 times with 1× BB. Finally, 400 μg of nuclear proteins (adjusted to a concentration of 1.6 mg/ml with nuclear lysis buffer) and an equal volume of buffer D (20 mM HEPES, pH 7.9, 100 mM KCl, 0.2 mM EDTA, 8% glycerol, 1x cOmplete™ Protease Inhibitor Cocktail (Roche), and phosphatase inhibitors) were added and incubated with rotation at 4 °C overnight. The next day, the beads were washed 3 times with 1× PBS, once with 100 mM NaCl and once with 200 mM NaCl. Bound DNA and proteins were eluted with 16 mM biotin (Sigma) in water (at pH 7.0) at 80 °C for 5 min, the eluate was transferred to a new tube and snap-frozen in liquid nitrogen.

Mass-spectrometric analysis of the eluates was performed with nano-LC-MS/MS using Q Exactive Plus (Thermo Scientific) at Proteomics core facility at the University of Tartu, Estonia, as described previously (Mutso et al., 2018) using label-free quantification instead of SILAC and Rattus norvegicus reference proteome for analysis. The full lists of proteins obtained from mass-spectrometric analysis are shown in Supplementary Table 3. Custom R-script was used to keep only transcription factors based on gene symbols of mammalian genes from gene ontology categories “RNA polymerase II cis-regulatory region sequence-specific DNA binding” and “DNA-binding transcription factor activity” obtained from www.geneontology.org (16.03.2020). At least 1.45-fold enrichment to the +3 kb enhancer probe compared to the +11 kb intronic probe was used as a cutoff for specific binding. The obtained lists were manually curated to generate Venn diagram illustration of the experiment.

### 8. Chromatin immunoprecipitation

Chromatin immunoprecipitation (ChIP) assay was performed as described previously (Esvald et al., 2020) using 10 min fixation with 1% formaldehyde. 5 μg of CREB antibody (catalog #06-863, lot 2446851, Merck Millipore) or TCF4 antibody (CeMines) was used per immunoprecipitation (IP). DNA enrichment was measured using qPCR. All qPCR reactions were performed in 10 μl volume in triplicates with 1× LightCycler 480 SYBR Green I Master kit (Roche) and primers listed in Supplementary Table 1 on LightCycler^®^ 480 PCR instrument II (Roche). Primer efficiencies were determined by serial dilutions of input samples and were used for analysing the results. Percent of input enrichments were calculated for each region and IP, and data were log-transformed before statistical analysis.

ENCODE data of different ChIP-seq experiments was visualized using UCSC Genome Browser track “Transcription Factor ChIP-seq Peaks (340 factors in 129 cell types) from ENCODE 3 Data version: ENCODE 3 Nov 2018”. Data of previously published ChIP-seq experiments were obtained from Gene Expression Omnibus with accession numbers GSM530173, GSM530174, GSM530182, GSM530183 (Kim et al., 2010), GSM1467429, GSM1467434 (Malik et al., 2014), GSM1820990 (Moen et al., 2017), GSM1647867 (Sun et al., 2019), GSM1649148 (Bayam et al., 2015), and visualized using Integrative Genomics Viewer version 2.8.0 (Robinson et al., 2011).

### 9. Statistical analysis

All statistical tests and tested hypotheses were decided before performing the experiments. As ANOVA’s requirement of homoscedasticity was not met, two-tailed paired or unpaired equal variance t-test, as reported at each figure, was used for statistical analysis using Excel 365 (Microsoft). To preserve statistical power, p-values were not corrected for multiple comparisons as recommended by (Feise, 2002; Rothman, 1990; Streiner and Norman, 2011).

## ACKNOWLEDGEMENTS

This work was supported by Estonian Research Council (institutional research funding IUT19-18 and grant PRG805), Norwegian Financial Mechanism (Grant EMP128), European Union through the European Regional Development Fund (Project No. 2014-2020.4.01.15-0012), H2020-MSCA-RISE-2016 (Grant EU734791), the Biotechnology and Biological Sciences Research Council (BB/M001199/1, BB/M007103/1, and BB/R001049/1). This work has also been partially supported by “TUT Institutional Development Program for 2016-2022” Graduate School in Clinical medicine receiving funding from the European Regional Development Fund under program ASTRA 2014-2020.4.01.16-0032 in Estonia. We thank Epp Väli and Andra Moistus for technical assistance, Indrek Koppel and Priit Pruunsild for critical reading of the manuscript.

## SUPPLEMENTARY FIGURES

**Supplementary figure 1.**
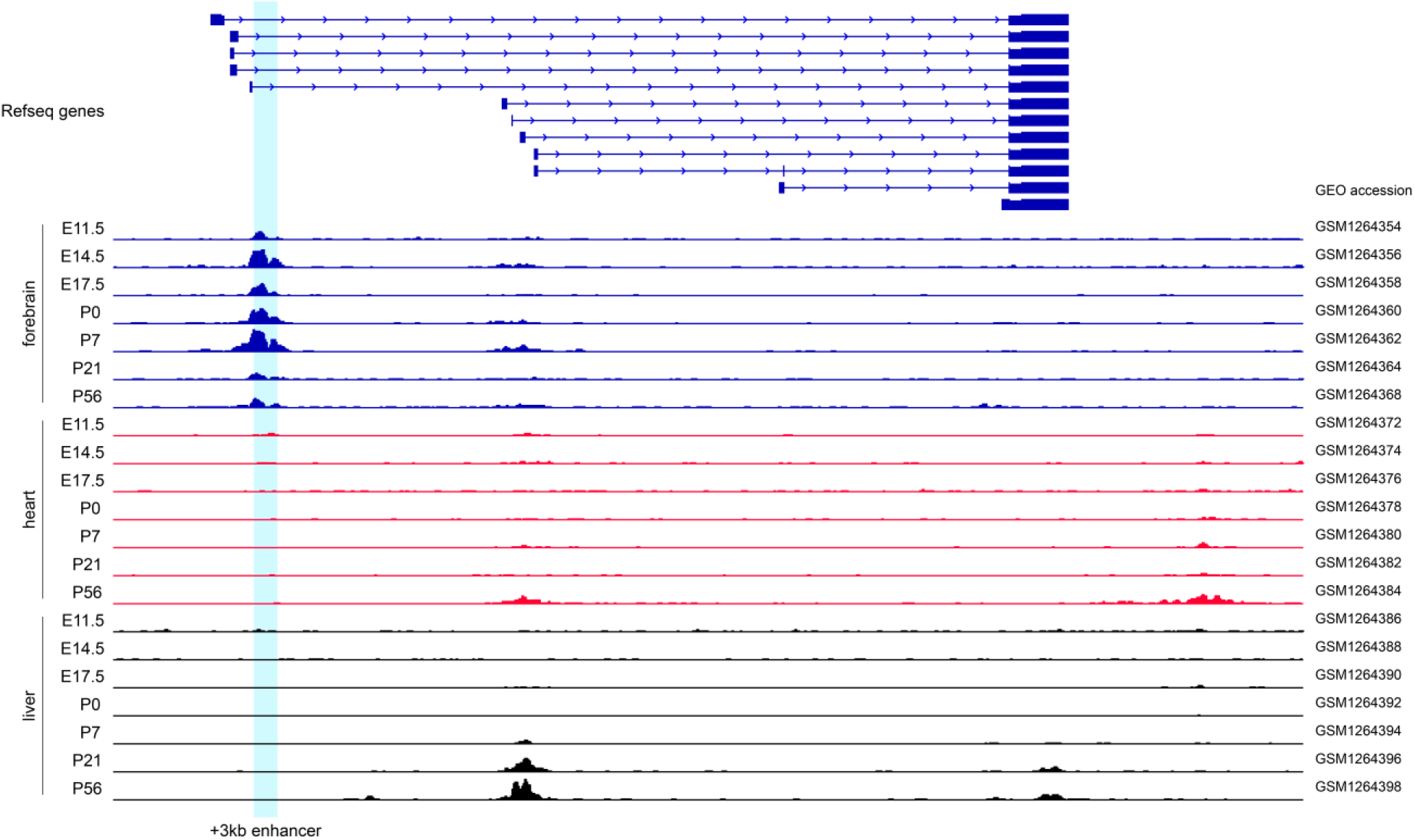
+3 kb enhancer region shows brain-specific H3K27ac histone modification. Integrative Genomics Viewer was used to visualize H3K27ac ChIP-seq data of different mouse tissues throughout the development (Nord et al., 2013). Different BDNF transcripts are shown in the upper part of the figure, developmental stage and tissue are shown on the left. E indicates embryonic day, P postnatal day. +3 kb enhancer region is marked with light blue. GEO accession numbers of the data are shown on the right.

**Supplementary figure 2.**
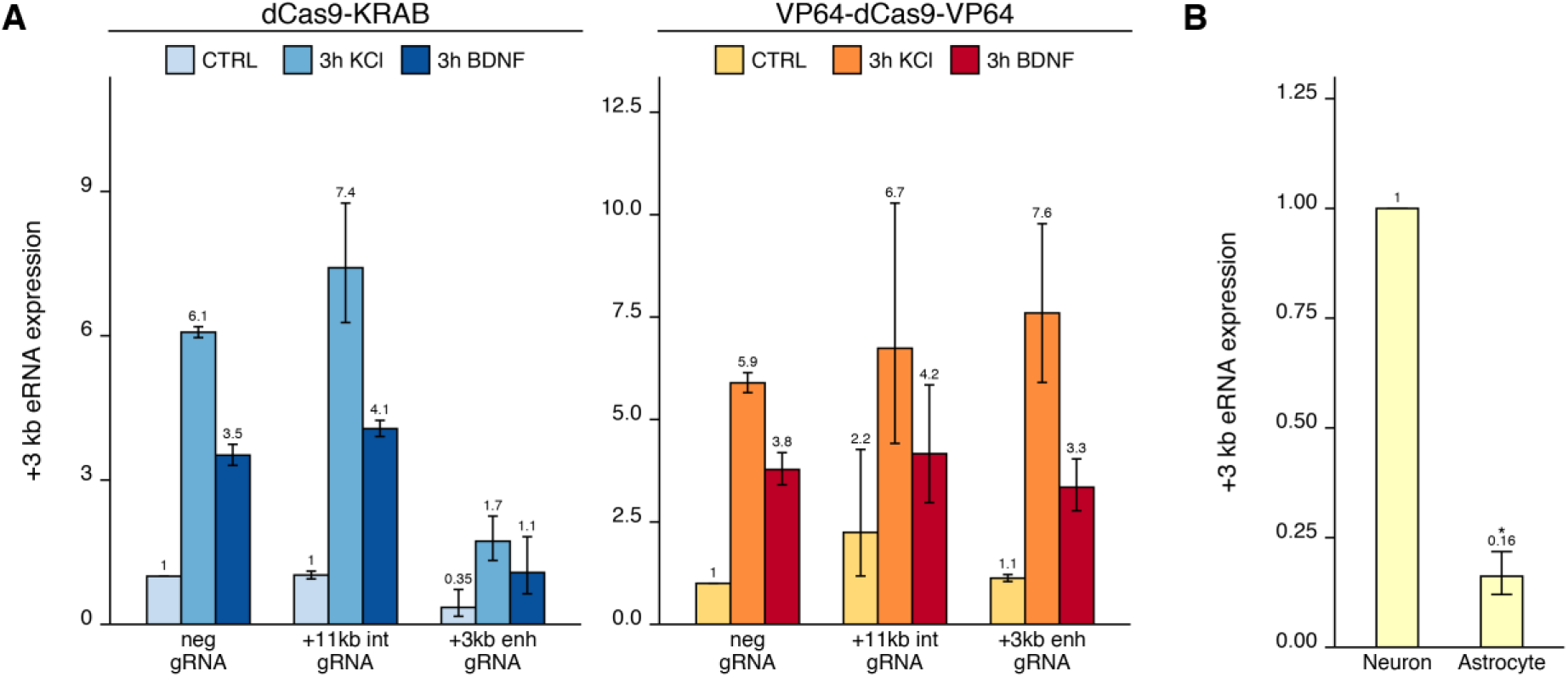
+3 kb enhancer shows stimulus-dependent eRNA transcription in neurons. (A) Measurement of +3 kb eRNA in CRISPRi and CRISPRa experiments in cultured neurons. Rat cultured cortical neurons were transduced at 0 DIV with lentiviral particles encoding either dCas9 fused with Krüppel associated box domain (dCas9-KRAB, blue) or 8 copies of VP16 domain (VP64-dCas9-VP64, orange) together with lentiviruses encoding either guide RNA that has no corresponding target sequence in the rat genome (neg gRNA), a mixture of four gRNAs directed to the putative +3 kb BDNF enhancer (+3 kb enh gRNA) or a mixture of four guide RNAs directed to +11 kb intronic region (+11 kb int gRNA). Transduced neurons were left untreated (CTRL) or treated with 50 ng/ml BDNF or 25 mM KCl (with 5 μM D-APV) for 3 hours at 8 DIV. Expression levels of +3 kb eRNA were measured with RT-qPCR and normalized to HPRT1 expression levels. eRNA expression levels are depicted relative to the eRNA expression in untreated (CTRL) neurons transduced with negative guide RNA within each set (dCas9-KRAB or VP64-dCas9-VP64). The average eRNA expression of independent experiments is depicted above the columns. Error bars represent SEM (n = 2 independent experiments). No statistical analysis was performed. (B) Comparison of +3 kb enhancer eRNA expression levels in cultured neurons and astrocytes. eRNA expression was measured using RT-qPCR, unnormalized average Ct values were used for the analysis and transformed to linear scale for graphical representation. The +3 kb eRNA expression level in neurons was set as 1. The average eRNA expression of independent experiments is depicted above the columns. Error bards represent SEM (n = 3 independent experiments). * p<0.05 (paired two-tailed t-test).

**Supplementary figure 3.**
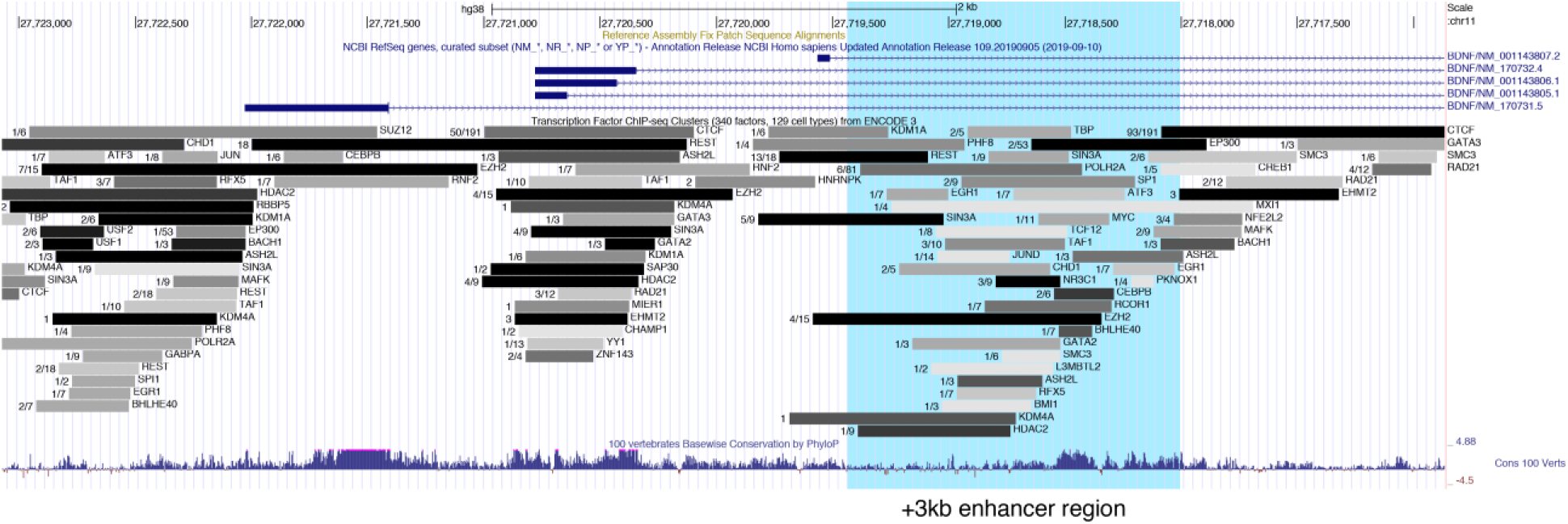
+3 kb enhancer region binds various transcription factors in human cell lines. UCSC Genome Browser was used to visualize ENCODE ChIP-seq data track “Transcription Factor ChIP-seq Peaks (340 factors in 129 cell types) from ENCODE 3 Data version: ENCODE 3 Nov 2018” at the +3 kb enhancer region. Numbers indicate cell lines with binding of the indicated transcription factor/number cell lines assayed for the binding of indicated transcription factor. The +3 kb enhancer region is shown in light blue.

